# Intrinsic functional connectivity delineates transmodal language functions

**DOI:** 10.1101/2024.12.20.629770

**Authors:** J.J. Salvo, N.L. Anderson, R.M. Braga

**Author notes:** Corresponding authors: Joseph J. Salvo, Rodrigo M. Braga.

## Abstract

Communication involves the translation of sensory information (e.g., heard words) into abstract concepts according to abstract rules (e.g., the meaning of those words). Accordingly, using language involves an interplay between *unimodal* brain areas that process sensory information and *transmodal* areas that respond to linguistic input regardless of the input modality (e.g., reading sentences vs. listening to speech). Previous work has shown that intrinsic functional connectivity (iFC), when performed within individuals, can delineate a distributed language network that overlaps in detail with regions activated by a reading task. The network was widely distributed across multiple brain regions, recapitulating an organization that is characteristic of association cortex, and which suggests that the distributed language network serves transmodal, not unimodal, functions. Here, we tested whether the distributed language network encapsulates transmodal functions by assessing its degree of overlap with two language tasks, one auditory (i.e., listening to speech) and one visual (i.e., reading sentences). The results show that the distributed language network aligns well with regions activated by both tasks, supporting a transmodal function. Further, the boundaries of the distributed language network along the lateral temporal cortex serve as a good proxy for the division between transmodal language and auditory functions: presentation of sounds (i.e., filtered, incomprehensible speech) evoked activity that was largely outside of the distributed language network but closely followed the network boundaries. These findings support that individualized iFC estimates can delineate the division between sensory-linked and abstract linguistic functions. We conclude that within-individual iFC may be viable for language mapping in individuals with aphasia who cannot perform language tasks in the scanner.

## 1 Introduction

Language allows us to communicate the same idea through multiple senses, including hearing (speech), vision (reading), or touch (braille). In each case, sensory percepts (i.e., words) are translated into their associated meaning according to linguistic rules. Thus, communication requires the coordination of a broad array of cognitive processes, including *unimodal* functions that process information from a given sense (e.g., visual, auditory, tactile) but also *transmodal* functions that transcend a particular sensory modality. These transmodal functions may include syntactic and lexical processes that are likely shared across different forms of language input. Accordingly, brain mapping studies that use auditory or visual linguistic stimuli have often reported overlapping activity (e.g., Scott et al., 2017; Fedorenko et al., 2010; Binder et al., 1997). A recent proposal unites these shared regions as a “core language network” that is sensitive to linguistic stimuli regardless of input modality (Fedorenko et al., 2024; see also Fedorenko & Thompson-Schill, 2014), but which interacts with modality-specific regions that provide sensory input and enable motor output. In this framework, the core language network sits in between unimodal sensory regions and higher-level regions that encode the meaning of words across association cortex (Huth et al. 2016).

Recent improvements in functional magnetic resonance imaging (fMRI) have allowed estimation of brain networks within individuals through repeated sampling (Laumann et al. 2015; Braga & Buckner 2017; Gordon et al. 2017). The within-individual estimates allow better separation of networks which are blurred together by a group-wise approach that averages data across individuals (Braga & Buckner 2017). We showed that within-individual intrinsic functional connectivity (iFC; Biswal et al., 1995) can delineate a distributed language network (LANG) from resting-state data, despite the absence of a language task (Braga et al., 2020; see also Glasser et al. 2016; Lee et al. 2012; Hacker et al. 2013; Branco et al., 2020). The LANG network included prominent regions within the inferior frontal cortex, at or near the classic Broca’s area, and in the lateral temporal cortex, at or near the classic Wernicke’s area. In addition, we noted that the network also included regions in the supplementary motor area (SMA), the temporopolar region, and possibly ventral temporal cortex (Braga et al., 2020; Nobre & McCarthy, 1995; Thomas et al., 2023, Li & Saygin, 2024). Confirming that this network serves linguistic functions, we showed that the full set of regions activated more strongly when participants read written sentences compared to lists of pronounceable, made-up words (i.e., pseudowords). This suggested that the whole distributed network was engaged during the processing of written sentences, and further supported that the network remains connected even in the absence of an overt language task.

An interesting observation was that when the full set of regions was considered, the organization of the LANG network recapitulated a parallel distributed network motif that is proposed to be characteristic of association cortex, Based on tracer studies of the macaque brain, Goldman-Rakic (1988) proposed that a parallel distributed network architecture, in which multiple association regions throughout frontal, parietal, temporal, and midline cortices are reciprocally interconnected, supports associative functions through brain-wide interactions. This architecture is fundamentally different from the stricter hierarchically-connected networks that serve unimodal sensory and motor functions (Felleman & Van Essen, 1991; Mesulam, 1998). The distributed nature of LANG therefore suggests that the network serves transmodal, rather than unimodal, language functions.

It is not known the degree to which this is true. Our previous study (Braga et al. 2020) used only visual stimuli (from Fedorenko et al. 2010; Fedorenko & Thompson-Schill 2014) which aimed to isolate higher-level aspects of language related to lexical (i.e., linking sensory percepts with word-level meanings) and combinatorial processes (i.e., linking multiple words into phrase- and sentence-level meanings), while controlling for sensory processes (i.e., reading combinations of letter strings that did not have word- and sentence-level meaning). Interestingly, the visual language localizer activated lateral temporal regions that encircled primary auditory areas and Heschl’s gyrus, despite the absence of auditory input (Braga et al. 2020; Fedorenko et al. 2010). This implies that these lateral temporal LANG regions serve transmodal language processes despite their proximity to auditory processing hierarchies (Rauschecker & Scott 2009).

Here, we sought to compare language-evoked activity across different sensory modalities to establish the degree to which iFC estimates of the LANG network represent transmodal language functions. We performed network estimation within individuals using iFC, then assessed whether the iFC-defined LANG network overlapped with activity evoked by visual (i.e., reading sentences; Fedorenko et al. 2010) and auditory language tasks (i.e., listening to speech; Scott et al., 2017). We also assessed how the LANG network boundaries related to regions active during auditory processing of incomprehensible speech sounds.

## 2 Materials & Methods

### 2.1 Subjects and sessions

Ten adults (6 female, ages 22-36, mean age 26.6, 9 right-handed) from the local community were recruited as part of the ‘Detailed Brain Network Organization’ (DBNO) study. Details regarding this dataset have been previously described in Kwon et al., 2024 and Edmonds et al., 2024. Participants had normal hearing and normal or corrected to normal vision, and no history of neurological illness. Informed consent was obtained from all participants for being included in the study. All participants provided written consent and were compensated for participation. Procedures were approved by Northwestern University’s Institutional Review Board. Participants were invited to 8 magnetic resonance imaging (MRI) sessions, each of which featured several cognitive tasks as well as a passive fixation (“resting-state”) task for network mapping using iFC. Prior to the first MRI session, participants were trained on all the tasks and were extensively coached about strategies for staying still in the scanner to improve data quality. While in the scanner, participants’ heads were padded using inflatable cushions to restrict head motion. Participants were also informed that the initial 1-2 MRI sessions would serve as a trial to assess compliance and whether they wanted to continue participation. Based on these criteria, 2 subjects were not invited back for additional sessions, leaving 8 subjects who completed all 8 MRI sessions (4 female, ages 22-36, mean age 26.75, 7 right-handed). This led to a total of 60.8 hours of fMRI data collected (7.6 hours per participant), for precision functional mapping.

### 2.2 Task Descriptions

#### 2.2.1 Passive fixation task (REST)

To allow estimation of large-scale networks within each individual using iFC, participants completed a passive fixation task in the scanner. Participants were shown a crosshair in the center of the screen and instructed to fixate on the crosshair for the duration of the task, keeping their eyes open and blinking normally. The task lasted ∼7 minutes, and each participant completed two runs in each session, as the first and final runs of the session (total: 16 runs, ∼112 minutes per participant). One participant (S2) completed an additional REST run to replace a poor-quality run that had been excluded in a prior session (see *MRI quality control*).

#### 2.2.2 Written language localizer (READLOC)

This task used visual stimuli (text) and lasted 5 minutes per run (from Fedorenko et al., 2010). In each trial, participants read a 12-word sentence or 12-pseudoword sequence on the screen, one word/pseudoword at a time. Participants were visually cued to press a button following each trial. Sentences were interleaved with sequences and presented in alternating blocks of three (1 block = 3 trials = 18.1s), with 36 trials per run (18 sentences, 18 pseudoword). A crosshair was presented briefly between trials, as well as after every four blocks for a period comparable length to the block duration (17.5s). Participants were visually cued by a hand icon to press a button with their right index finger following each sequence, as a means of maintaining alertness.

#### 2.2.3 Auditory language localizer (SPEECHLOC)

This task used auditory stimuli (speech) and lasted 7 minutes per run (based on Scott et al., 2017). In each trial, participants listened to audio clips of normal intelligible speech and unintelligible, distorted speech. Audio clips were taken from recordings of TED talks, The Moth Radio Hour, and Librivox audiobooks. Two clips were taken from each recording, with one clip left intact and the second distorted by filtering as outlined below. This meant that the distorted clips preserved certain sound characteristics (speaker identity, intonation, speed, recording quality, etc.) to improve their effectiveness as control stimuli.

Each clip was edited to 18 seconds in duration, and was volume standardized and frequency equalized. All clips were compressed using a threshold of -40dB, and were transformed such that their root-mean-square equaled 0.0628. These values were chosen based on qualitative assessments of sound quality and volume by experimenter J.S. Stereo clips were converted to mono (same output for both ears). Clips were equalized using earphone-specific filters provided by the earphone manufacturer, Sensimetrics (Gloucester, USA). Without this filtering, clips sounded unnaturally shifted toward higher frequencies (“tinny”). We ultimately used the left-ear filter for both channels after pilot participants reported improved sound clarity.

For the distorted speech condition, half of the clips were filtered and combined with white noise to remove comprehensible speech, following Scott et al. (2017). To prevent filtering artifacts, dummy vectors 1/10th the length of each sound clip, with a magnitude matching the clip’s mean dB, were added to the start and end of each clip. Clips were lowpass filtered using a Butterworth infinite impulse response filter (IIR: Butterworth), with a pass frequency (Fpass) of 500 dB, a stop frequency (Fstop) of 600 dB, a passband ripple (Apass) of 1 dB, and a stopband attenuation (Astop) of 146 dB. White noise was generated by shuffling time points in the original clip and saving the resulting noise to a new vector. This vector was then transformed according to the amplitude envelope of the original clip, such that the volume of the noise was modulated by the volume of the voice. The noise vector was also lowpass filtered (IIR: Butterworth, Fpass=7500 dB, Fstop=10700 dB, Apass=1 dB, Astop=40 dB). Noise and voice were combined with a volume ratio of 1:2. The combined clip was lowpass filtered once more, using a threshold of 8000 dB, to remove noise-related spikes. Finally, the clip was trimmed back to a standard duration of 18s. To avoid clicking artifacts at the start and end of each sample, both intact and distorted clip volumes were ramped according to a y=x or y=1-x slope, applied respectively to the clip’s first and last 0.01 seconds.

Importantly, although the distorted clips were incomprehensible (i.e., participants could not make out the actual words being spoken), the clips preserved certain properties of speech, such as prosody and intonation.

Audio was presented through Sensimetrics S14 MRI-compatible earphones. To ensure that the clips were audible over the scanner noise, at the start of each MRI session, prior to data collection, we tested sound volume while running a dummy version of the MR imaging sequence to mimic the conditions of the task. Participants were played two speech clips (one clear, one distorted) followed by the tone used as a button-press cue. Participants used the provided button box to adjust the volume to a comfortable level at which the speech could be clearly heard, and were asked to describe the topic of the clearly-presented story.

Audio clips lasted 18 seconds and were followed by a tone lasting 0.15 seconds. Participants then had 2 seconds to press a button with their right index finger to register attention. Each task run included 8 clear and 8 distorted 18s audio clips. Four clips were presented as a block (two clear, two distorted), with four blocks per run. A fixation cross (‘+’) was presented between blocks for 14 seconds. Within each block, clips were presented in a counterbalanced order, e.g., CDDC+DCDC+CDCD+DCCD (C: clear clip, D: distorted clip, +: fixation), ensuring an even distribution of conditions across the run. Block order was also counterbalanced across runs.

Participants completed additional tasks in each session that are not described in this report (see Edmonds et al., 2024 and Kwon et al., 2024).

### 2.3 MRI data acquisition

Data were collected on a Siemens 3.0 T Prisma scanner (Siemens, Erlangen, Germany) at the Center for Translational Imaging at Northwestern University’s Feinberg School of Medicine in Chicago. Two anatomical images were collected: a T1-weighted image in the first MRI session (TR = 2,100ms, TE = 2.9ms, FOV = 256 mm, flip angle = 8°, slice thickness = 1 mm, 176 sagittal slices parallel to the AC-PC line) and a T2-weighted scan in the second session (TR = 3,000ms, TE = 565ms, FOV = 224 mm x 256 mm, flip angle = 120°, slice thickness = 1 mm). Both scans included volumetric navigators (Tisdall et al., 2012) from the Adolescent Brain Cognitive Development study (Hagler et al., 2019). Functional MRI data were collected using a 64-channel head coil using a multi-band, multi-echo sequence with the following parameters: TR = 1,355ms, TE = 12.80ms, 32.39ms, 51.98ms, 71.57ms, 91.16ms, flip angle. = 64°, voxel size = 2.4mm, FOV = 216 mm x 216 mm, slice thickness = 2.4 mm, multiband slice acceleration factor = 6; see Poser et al., 2006; Lynch et al., 2020).

### 2.4 Quality control

Head motion was estimated using FSL’s MCFLIRT (Jenkinson et al., 2002). Runs were automatically excluded from analysis if motion exceeded predetermined thresholds of framewise displacement (FD) > 0.4mm or absolute displacement (AD) > 2.0mm. Runs were flagged for visual inspection if motion exceeded more stringent thresholds (FD > 0.2mm, AD > 1.0mm), and the whole run was then excluded if motion could be clearly seen in the raw data. Runs for the remaining three localizer tasks could also be excluded due to poor behavioral performance. For READLOC, runs with more than 5% missing button-presses were excluded. For SPEECHLOC, runs with more than 20% missing button-presses were excluded (as there were fewer total trials).

After quality control, each participant retained at least 10 REST runs (range: 10-16). For all participants except one (S5), half of the data were set aside for replication analyses (not used in this study), leaving 49-60 minutes of data for each participant (S1: 56 min; S2: 56; S3: 56; S4: 56; S5: 60; S6: 49; S7: 49; S8: 49).

For SPEECHLOC, at least 7 runs were retained per subject (S1: 8; S2: 8; S3: 8; S4: 7; S5: 7; S6: 8; S7: 7; S8: 8) for a range of 49 to 56 minutes of data per participant. For READLOC, at least 4 runs were retained per subject (S1: 8; S2: 8; S3: 8; S4: 4; S5: 7; S6: 8; S7: 5; S8: 7) for a range of 20 to 40 minutes of data per participant.

### 2.5 MRI data preprocessing

Blood-oxygenation-level-dependent (BOLD) data were preprocessed using a custom processing pipeline “iProc” (Braga et al., 2020) optimized for within-individual alignment of data from multiple runs and sessions. When possible, smoothing and interpolations were combined in an effort to minimize blurring. Additional steps were included to account for the multi-echo data (outlined below). Each individual’s data were processed separately. The first 9 volumes (∼12 s) of each run were discarded to remove a T1 attenuation artifact. Next, a mean BOLD template was created as an interim stage for data registration by averaging the first echo of all included runs of all tasks, to reduce bias towards any one particular run. The participants’ T1 anatomical images were used to create a native space template. These templates were used to build four matrix transforms to align each BOLD volume 1) to the middle volume of the same run for motion correction, 2) to the mean BOLD template for cross-run alignment, 3) to the native space template, and 4) to the 152 1-mm atlas (Mazziotta et al. 1995) from the Montreal Neurological Institute (MNI) International Consortium for Brain Mapping (ICBM). The four transforms were composed into two matrices, which were then applied to the original volumes to register all volumes to the T1 anatomical template (matrices 1-3) and to the MNI atlas (matrices 1-4) in a single step. The MNI registration was used to regress non-gray matter signals from resting state runs (*see Functional connectivity preprocessing* below). Visual checks were incorporated into each registration step of the pipeline.

Motion correction transforms were calculated using rigid-body transformation based on the first echo, which had less signal dropout and better preserved the shape of the brain. Registration matrices were calculated based on the first echo, and then applied to all the echoes. Once data were projected to the T1 native space, the five echoes were combined to approximate local T2* by weighting each echo according to its temporal signal-to-noise (tSNR) ratio and echo time (Heunis et al., 2021). Briefly, the tSNR was calculated for each echo, which was then weighted by the echo time. This weighted tSNR is then divided by the sum of all weighted tSNRs (i.e., for all echoes). The resulting image was multiplied by the echo’s original intensity image, and these were summed across all echoes to create the final optimally-combined image.

### 2.6 Functional connectivity preprocessing

Nuisance variables were calculated for each run, by extracting signal averages from deep white matter and from cerebrospinal fluid using masks hand-drawn in MNI space that were back-projected to the native space. Additional nuisance variables included 6 motion parameters, whole-brain (global) signal, and their respective temporal derivatives. Data were then bandpass filtered using a range of 0.01–0.1 Hz using *3dBandpass* (AFNI v2016.09.04.1341; Cox, 1996; 2012). Next, data were projected to a standard cortical surface (fsaverage6, 40,962 vertices per hemisphere; Fischl, 1999), and smoothed with a 2mm full-width at half-maximum kernel along the surface. This kernel size was chosen based on prior work (Braga & Buckner, 2017; Braga et al., 2019) to retain functional anatomic detail while limiting noise (i.e., speckling) in the functional connectivity maps.

### 2.7 Network estimation

Prior to any analysis of the task data, the independent REST data were used to create subject-specific estimates of large-scale networks using iFC following our previously used procedures (Braga & Buckner, 2017; Braga et al., 2020). First, functional connectivity matrices were calculated for each REST run by computing vertex-vertex Pearson’s product-moment correlations. The matrices were then z-normalized using the Fisher transform, averaged across all runs within each individual, then converted back to r values using the inverse Fisher transform. These within-individual average matrices were used to perform a seed-based analysis. Seeds were manually selected in the left hemisphere to delineate 8 networks, including LANG; see Braga & Buckner, 2017; Braga et al., 2020; Edmonds et al., 2024; Kwon et al., 2024) and in the right hemisphere to identify the auditory network, AUD (to ensure the left-hemisphere correlated regions observed were not due to local autocorrelation effects). We then used a multi-session hierarchical Bayesian Model (MS-HBM) approach to define networks using a data driven algorithm applied to the same data (Kong et al., 2019). This method stabilizes individual-specific network estimates by integrating priors from multiple levels (e.g., group atlas, cross-individual and cross-run variation). For each individual, we generated clustering solutions with k values (i.e., number of clusters) between 12–18, and selected the lowest level of clustering that separated the networks of interest as defined by the seed-based approach. The MS-HBM algorithm also allowed definition of networks beyond the *a priori* selected networks, including those covering auditory cortex (surrounding bilateral Heschl’s gyrus; AUD) and ventral somatomotor cortex (surrounding bilateral inferior central sulcus).

### 2.8 Task activation maps

Data from the localizer tasks were analyzed using FSL’s FEAT (Woolrich et al., 2001). Each hemisphere of each surface-projected task run was input into a separate general linear model (GLM). The task was modeled using explanatory variables that were convolved with a double-gamma hemodynamic response function using FEAT.

For SPEECHLOC, regressors were specified that included all 18-second presented audio clips for each of the two speech types (comprehensible and distorted). Beta values were calculated for the comprehensible and distorted speech conditions, and a contrast of parameter estimates was calculated between the conditions for each run. The contrast maps from each run were normalized into z values by FEAT, and then averaged across all runs for a given task and individual.

An additional contrast map, AUDLOC, was created focusing on the distorted speech condition of SPEECHLOC, contrasted implicitly against the fixation baseline periods. This was used to localize auditory cortex (i.e., regions active during sound presentation compared to baseline). Because the distorted speech condition preserved some features of speech sounds, such as prosody and intonation, this contrast also likely activates speech-selective regions sensitive to these features that may lie outside primary and secondary auditory cortex.

For READLOC, regressors were specified that included all 18.1-second blocks of reading for each of the two block types (real words and pseudowords). Beta values were calculated for the real words and pseudowords conditions, and a contrast of parameter estimates was calculated between these conditions for each run. The contrast maps were normalized into z values by FEAT, and then averaged across runs for a given task and individual.

To ease visualization of overlap, we thresholded and binarized the maps from each of the task contrasts. First, we overlaid the maps from the two active language localizers (READLOC, SPEECHLOC) to identify regions responding to linguistic stimuli from both modalities. Second, we overlaid the maps from the READLOC and AUDLOC contrasts to test the degree of overlap between transmodal language and auditory regions with the iFC-estimated LANG network. The latter two contrasts were chosen so that the estimate of transmodal language regions (from READLOC) was from independent data to the maps of auditory regions (from AUDLOC), to ensure that any overlap or lack thereof was not due to the contrasted conditions.

In a final analysis, we quantified how closely the vertices that overlapped between these maps aligned with the LANG network in the left hemisphere. For each candidate vertex, we first determined whether the vertex fell within the boundaries of the LANG network (distance of 0). If not, all neighboring vertices on the surface mesh were identified (those sharing a face with the candidate vertex). If any of these fell within the LANG network, the distance for the candidate vertex was 1. If not, we identified all vertices sharing a face with these neighboring vertices (second-level neighbors), and so on. This process was repeated until a vertex within the LANG network was located, and a distance value was assigned to the candidate vertex. In essence, this determined the shortest distance between each candidate vertex and the LANG network.

## 3 Results

### 3.1 The distributed language network was defined using multiple intrinsic functional connectivity approaches

We defined large-scale networks *a priori* in each individual using iFC analysis of the independent passive fixation (i.e., “resting-state”) data. Seeds were chosen in lateral temporal cortex in the left hemisphere to identify the canonical left-lateralized language network (LANG). Seeds were also chosen in the right lateral temporal cortex to identify the primary auditory network (AUD) in the contralateral hemisphere (Fig. 1). Note that left-hemisphere seeds were also able to define the AUD network, despite being adjacent to the seed used to define LANG (not shown). As a complementary means to define networks, correlated vertices were clustered together using a data-driven approach (Fig. 1), with the number of clusters selected based on visual correspondence with the seed-based iFC maps, respecting that networks may be over-split or -lumped at different clustering levels. Candidate LANG and AUD networks were identified at a clustering solution of 14 clusters for subjects S1-S7. For S8, the candidate LANG network from the *k* = 14 solution, was adjacent to but did not align well with the seed-based maps of the LANG network nor the language-localizer maps. Therefore, for this participant, we *post hoc* changed the LANG and AUD estimates to be from the 15-cluster solution. Both analysis methods revealed a network comprising classic language regions, including in inferior frontal, lateral temporal, and posterior middle frontal cortices. Further, the results distinguished the LANG network from a neighboring network in lateral temporal cortex, AUD, that likely covers primary and secondary auditory areas.

**Figure 1:**
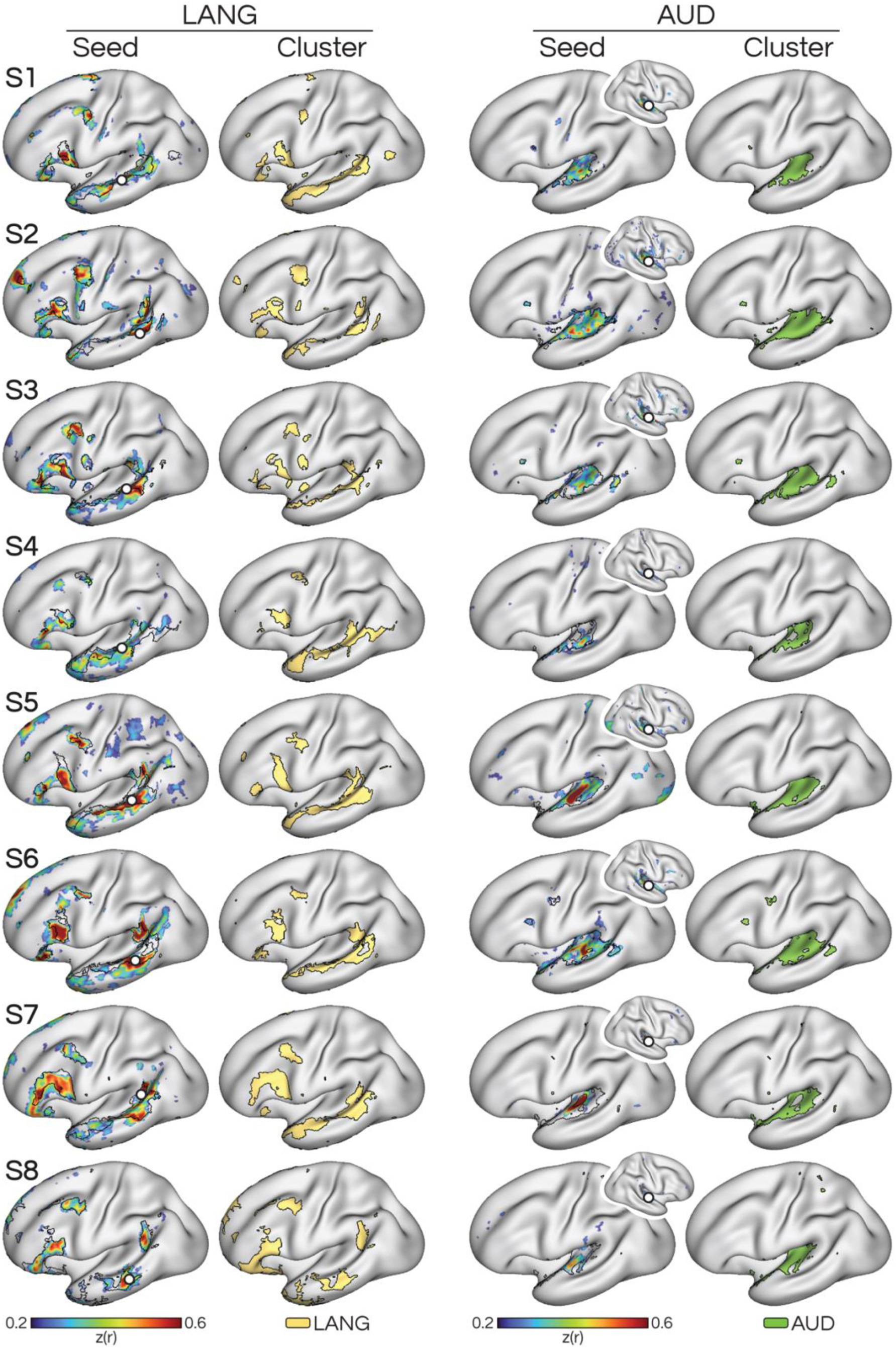
Within-individual functional connectivity estimates reveal separate, adjacent networks in superior temporal cortex. Left) Seeding in left lateral temporal cortex reveals a correlated network that includes regions classically associated with language processing (LANG; see Braga et al. 2020), including in inferior frontal cortex. The same network was also revealed by a data-driven clustering analysis, which provided converging estimates of the network (see how black boundaries of the clustering solution overlap with regions showing high correlation in the seed-based map). Right) Seeding in a nearby analogous location in right lateral temporal cortex reveals a different network (AUD) that included left hemisphere regions associated with auditory processing. This network was also confirmed as a distinct network by the clustering analysis. The two networks contained adjacent regions throughout the superior temporal cortex in the left hemisphere (see zoom-ins in Fig. 4).

### 3.2 Confirmation that the distributed language network is activated by a visual (reading-based) language task

We first replicated the results of Braga et al. (2020) by showing that the iFC-defined LANG network we selected supports linguistic functions. Analysis of the READLOC task showed greater activity within the LANG network while participants read comprehensible sentences, contrasted with reading sequences of pseudowords (Fig. 2, left). Notably, regions showing the greatest activity fell within the bounds of the LANG network, and in multiple regions the boundaries of the LANG network overlapped closely with activity evoked by the task. These results replicated that the LANG network is engaged at the network level during a reading task.

**Figure 2:**
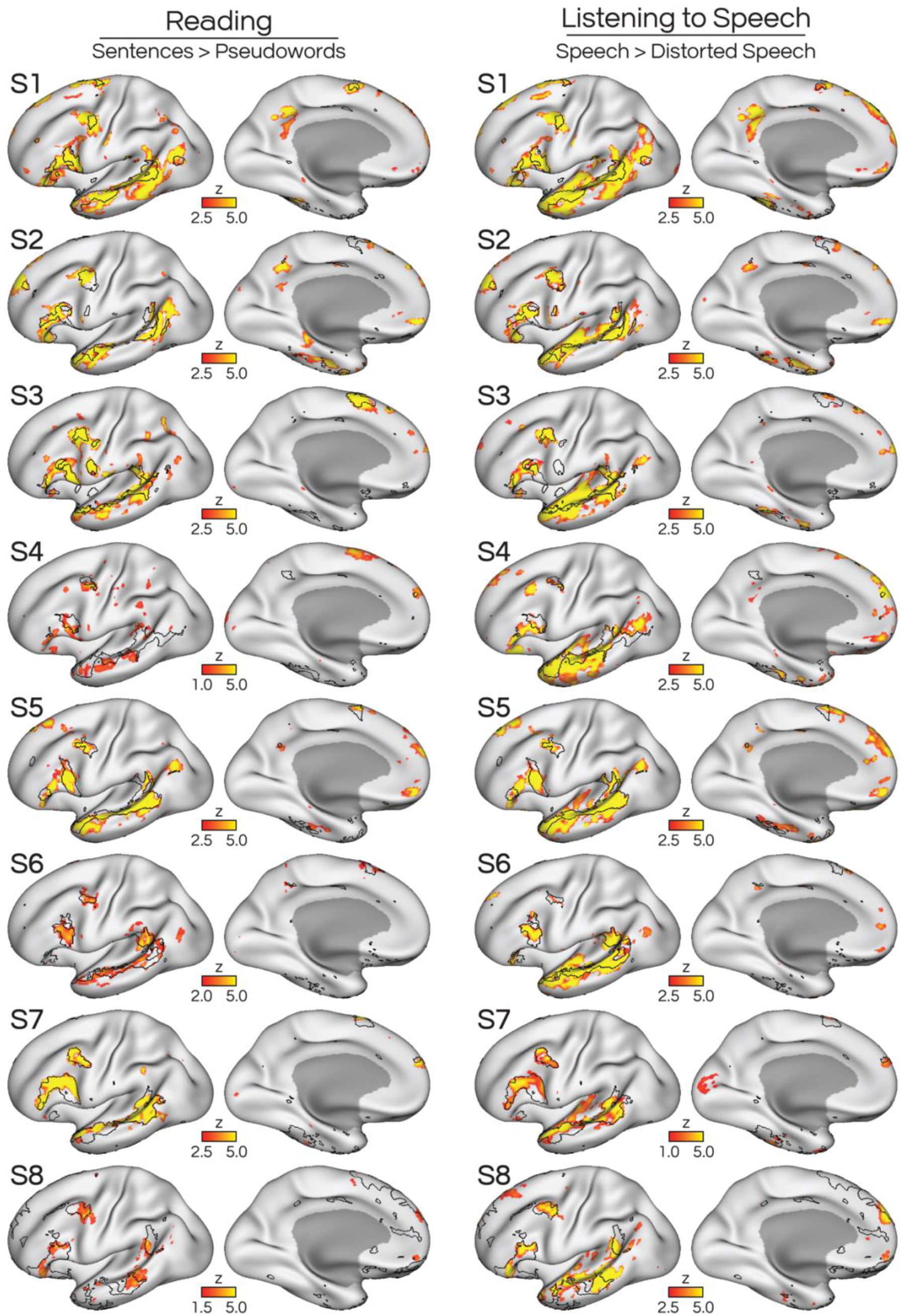
Visual language input (reading) and auditory language input (listening to speech) both robustly activated the distributed LANG network. Left) We replicated the results of Braga et al. (2020) in a new dataset, by showing that a reading task (READLOC; reading sentences contrasted against reading pseudoword sequences) elicited activity that overlapped with the iFC-derived estimate of LANG (see black borders from clustering solutions shown in Fig. 1). Detailed correspondence between iFC- and task-based estimates was seen in lateral temporal, inferior frontal, and posterior middle frontal cortex, as well as the pre-supplementary motor area (SMA). Thresholds were picked for each participant to highlight regions of strong activity while minimizing speckling (indicating noise). However, using the same threshold for all subjects revealed similar results. Note that for two subjects (S4, S8) the READLOC task did not elicit strong activation maps, so their thresholds were lowered to reveal a similar distribution of activated regions. Right) An auditory version of the language localizer task (SPEECHLOC; contrasting listening to sentences against listening to distorted speech) revealed a similar distribution of regions, which also overlapped with the FC-defined LANG network in multiple cortical locations. The SPEECHLOC map also included broader lateral temporal activity which likely extended into auditory regions, as explored in subsequent figures.

### 3.3 The distributed language network encompasses transmodal language functions

To establish whether the iFC-defined LANG network serves a transmodal language function, we analyzed the SPEECHLOC task data, which contrasted listening to clear, comprehensible speech with distorted, incomprehensible speech. Increased activity was observed for this contrast throughout the LANG network, including in the lateral superior temporal cortex, inferior frontal cortex, and posterior middle frontal cortex (Fig. 2, right). Some subjects (S1, S2, S3, S5) also showed activity in the SMA and anterior superior frontal cortex. Active regions consistently fell within the boundaries of the iFC-defined LANG network, often closely matching each subject’s unique functional anatomy (i.e., location and shape of network regions). We next created an overlap map between activation during reading (READLOC) and speech comprehension (SPEECHLOC) in each subject. Despite language input being from two different modalities, we observed extensive overlap of these two tasks within LANG network regions (Fig. 3, left). To quantify this, we took the surface vertices that showed overlap between the SPEECHLOC and READLOC contrasts, and calculated their distance to (i.e., number of vertices away from) the LANG network. Across all 8 subjects, the majority of vertices fell within the LANG network, and those outside were a short distance away (Fig. 3, right). This confirmed that the LANG network likely supports transmodal language processes, and likely encapsulates regions activated by linguistic input regardless of modality.

**Figure 3:**
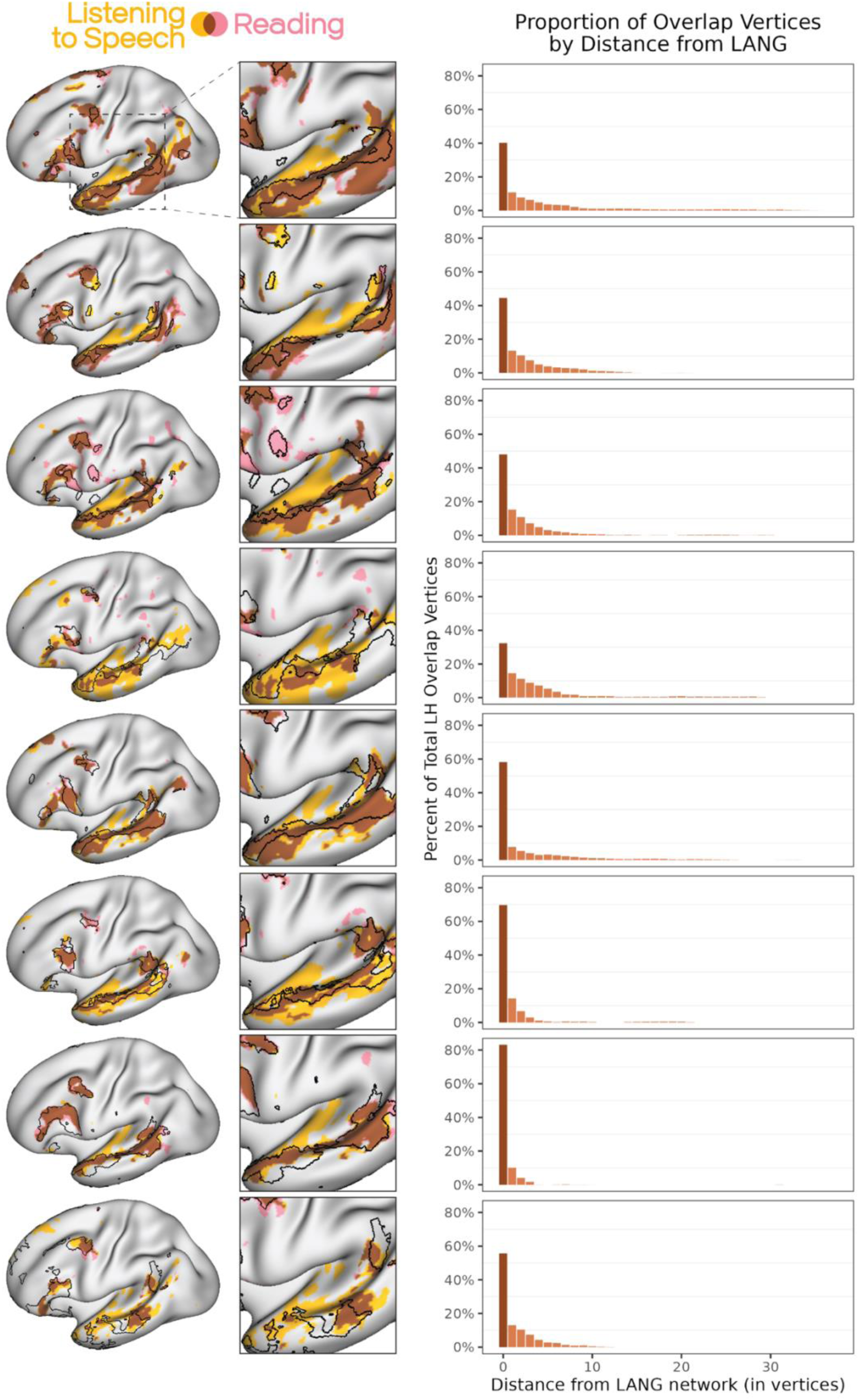
Overlap between activity elicited by reading and listening to speech is primarily within the distributed LANG network. Left) Regions activated by the auditory SPEECHLOC task overlapped extensively with those activated by the visual READLOC task. This overlap was predominantly within the bounds of the LANG network as defined with iFC, implying that the LANG network encapsulates transmodal language functions that engage with linguistic stimuli regardless of input modality. Right) Vertices showing overlap between the two task maps were quantified with respect to their proximity to the LANG network; in all subjects, the majority of overlap vertices are within the LANG network (a distance of 0) and the remainder were within a short distance of LANG, with the incidence of overlap vertices falling off with greater distance from the network.

For the SPEECHLOC contrast, we also saw activation that extended into the sylvian fissure that was not seen in READLOC, suggesting that the SPEECHLOC contrast was also revealing activity within unimodal auditory regions. These observations in combination raised the prospect that the boundaries of the iFC-defined LANG network might provide a relatively accurate delineation between modality-specific (i.e., auditory) and transmodal aspects of language processing.

### 3.4 Activity while listening to unintelligible speech activates primary auditory cortex

As a functional localizer for unimodal auditory regions, AUDLOC, we contrasted the distorted speech condition from the SPEECHLOC task with the fixation baseline of the same task. The resulting map revealed bilateral activation of superior lateral temporal areas that were at or near Heschl’s gyrus (Fig. 4), indicating that this was an effective contrast for isolating auditory cortex. Additional smaller regions of activity were also found in prefrontal cortex in some subjects (discussed later). Listening to distorted speech most extensively recruited regions in the AUD network, outside of the LANG network. However, we did observe some degree of overlap into the LANG network, to a varying amount across subjects.

**Figure 4:**
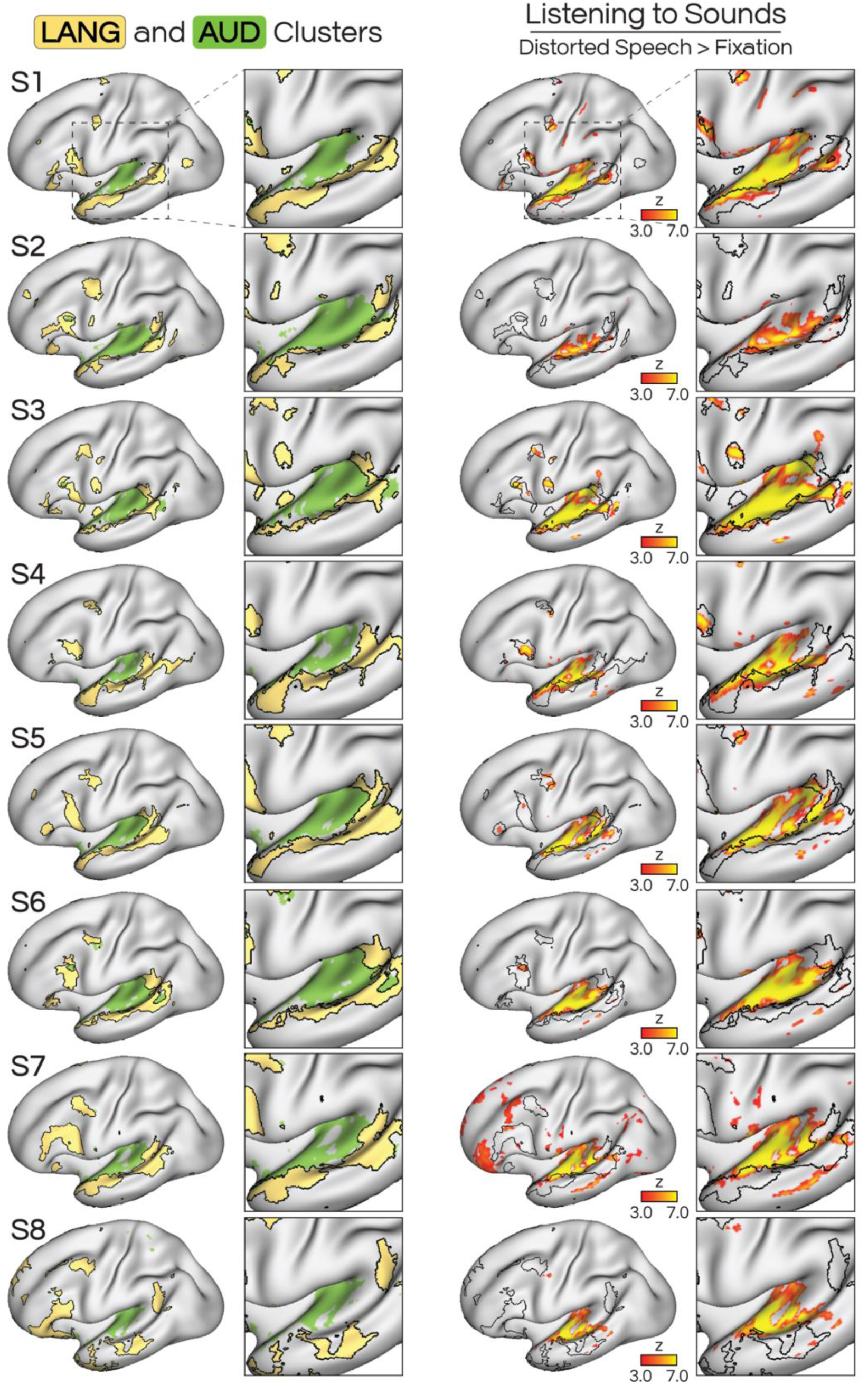
Auditory non-linguistic input (listening to distorted, incomprehensible speech) activates the AUD network, with limited activation of adjacent LANG network. Left) The iFC-defined AUD and LANG networks included adjacent regions, with LANG (yellow) encircling AUD (green) throughout the dorsal bank of the lateral temporal cortex. Right) The AUDLOC contrast contrasted listening to distorted speech against a passive fixation baseline as a localizer for auditory cortex. This map showed that listening to sounds revealed activity mainly within the AUD network, partly extending into the neighboring LANG network, however this was observed to varying degrees across subjects. We explored this overlap further in Fig. 5.

### 3.5 The boundaries of the distributed language network separate unimodal and transmodal functions

We explored the extent to which the iFC-defined LANG network separated transmodal and unimodal functions. To do so, we compared the thresholded, binarized maps from the READLOC contrast with the AUDLOC contrast (Fig. 5, left). First, we observed that the boundaries of the LANG network provided a remarkably good proxy for the demarcation of auditory non-linguistic from transmodal linguistic functions. In many cases, the areas of overlap between the maps were few, and the adjacent regions from each task were positioned close to, if not exactly at, the boundaries of the LANG network (see insets in Fig. 5). In some cases, the two maps overlapped, indicating that some regions that responded to the distorted speech stimuli were also recruited by the reading task. Notably, the regions that showed overlap between these maps were restricted almost entirely to the LANG network. To quantify this, we took the surface vertices that showed overlap between READLOC and AUDLOC, and calculated their distance to (i.e., number of vertices away from) the LANG network. Across 7 of the 8 subjects, the vast majority of the vertices fell within the LANG network, and those outside were a short distance away (Fig. 5, right). For S8, most vertices fell just outside the LANG network, possibly related to issues with network definition in this subject (see *Network estimation*). Nonetheless, this analysis revealed that iFC provides a remarkably accurate delineation between unimodal auditory and transmodal language functions along the left lateral temporal cortex.

**Figure 5:**
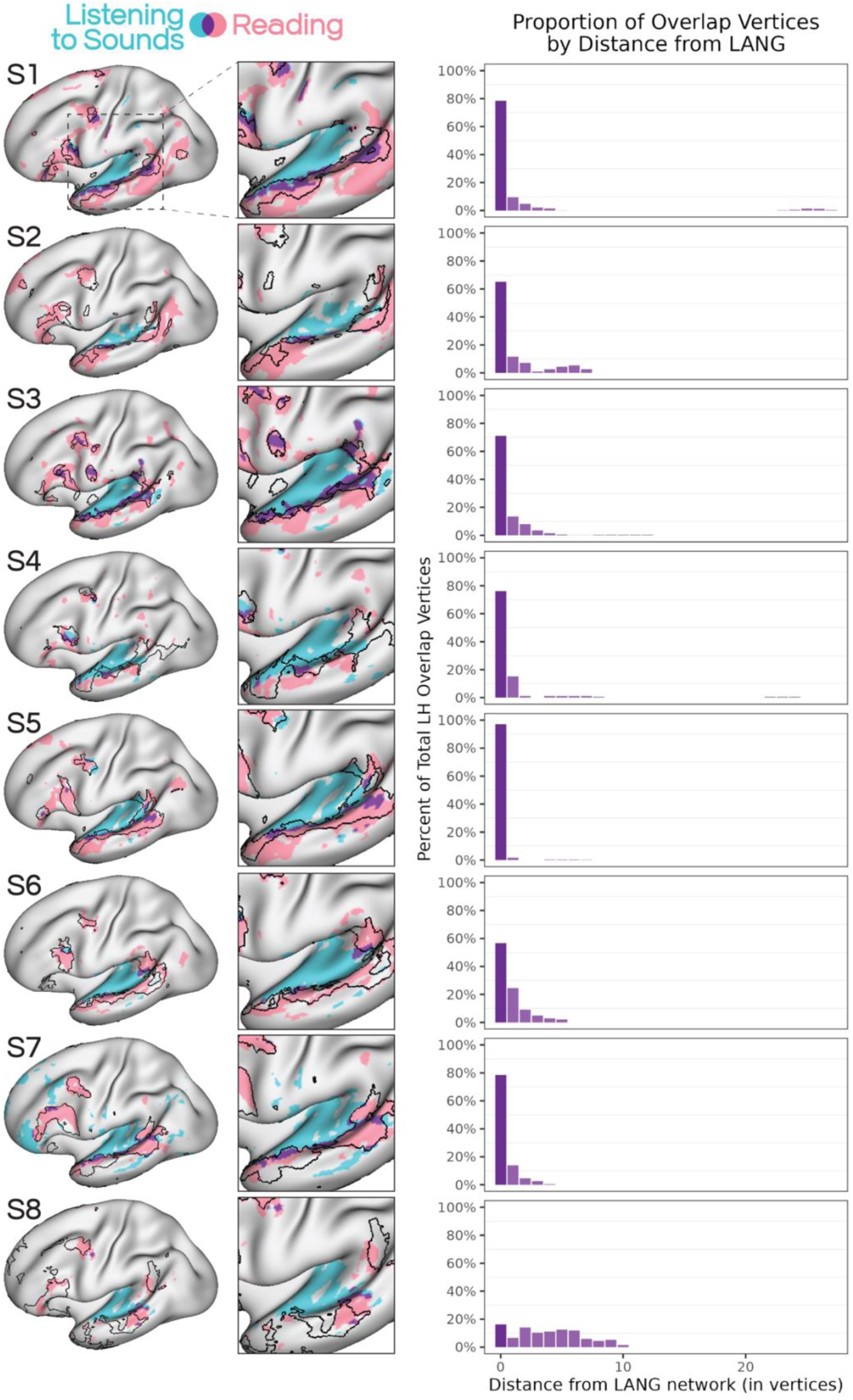
The boundaries of the LANG network in superior temporal cortex delineated between auditory and transmodal functions. Left) Regions that show increased activity during reading sentences (from READLOC; see Fig. 2), chosen here to represent transmodal language functions (see Fig. 3), were overlaid with regions activated by listening to non-linguistic sounds (the AUDLOC contrast; see Fig. 4). The insets show that the borders of LANG in many cases captured the transition between unimodal auditory and transmodal language functions. In other cases, overlap was sometimes seen between the two maps, however this overlap was mostly within the bounds of the LANG network. This overlap could be due to technical reasons (e.g., blurring) or the fact that the distorted speech stimuli preserved some features of speech, such as the intonation and prosody of the speaker, even though the spoken words were made incomprehensible. Right) Vertices showing overlap between the two maps were quantified with respect to their proximity to the LANG network; in subjects 1-7, the majority of overlap vertices were within the LANG network (a distance of 0), and the others were few vertices away.

## 4 Discussion

### 4.1 Overview

We investigated the properties of a distributed large-scale network that is associated with language processing (LANG). We used intrinsic functional connectivity (iFC) of resting-state data to define LANG as an intrinsically connected network that included classic language regions in inferior frontal (i.e., “Broca’s area”) and lateral temporal cortex (i.e., “Wernicke’s area”; Geschwind, 1965) in 8 individuals (Fig. 1). We then showed that the LANG network responds to linguistic stimuli presented both through visual (writing; Fig. 2) and auditory (speech; Fig. 3) input, and that the responses closely overlapped the network as estimated using iFC. The network included multiple regions along the superior temporal cortex that were adjacent to auditory cortex, but were clearly dissociated from unimodal auditory functions by an auditory localizer task contrast (AUDLOC, distorted speech clips > fixation, Fig. 4 & 5). Taken together, these findings advance our understanding of the language network as a transmodal system that works closely with adjacent, but functionally distinct, unimodal sensory areas.

### 4.2 The distributed language network serves a transmodal language function

Previous work has linked the iFC- defined LANG with regions active during reading (Braga et al., 2020), and other work has shown that a similar set of regions activate during both visual and auditory language stimuli (Fedorenko et al. 2010; Scott et al. 2017). Here, we show that auditory and visual language tasks both activate overlapping regions within each participant, and that the overlapping regions are well-encapsulated by the iFC-based estimate of the LANG network (Fig. 3). Thus, our results support that iFC is capable of mapping transmodal language functions within individuals.

This finding has high potential value, as language mapping is routinely performed for clinical purposes. For instance, mapping of language functions in individuals with language difficulties (i.e., aphasias) could inform prognoses by characterizing the extent of damage to language regions (Saur et al., 2010; Heiss & Kidwell, 2014; Wilson & Schneck 2021). Similarly, for patients with epilepsy or brain tumors, language is routinely used for surgical planning of resections (Tharin & Golby, 2007), guiding which areas may be removed while sparing language ability. A major limitation has been that some patients, particularly those with aphasia (but also those with other forms of dementia), may not be able to perform language tasks. Relatedly, task engagement may be hindered by scanning conditions (e.g., scanner noise during an auditory task, eyesight problems during a reading task), difficulties with task instructions, or participants failing to maintain attention to stimuli. These complications can prohibit conventional task-based language tasks from accurately activating language regions. Our current results indicate that iFC may provide a relatively accurate delineation of unimodal and transmodal language functions using only resting-state data, circumventing many of these potential obstacles.

### 4.3 The distributed language network demarcates transmodal and auditory functions

Both language localizer tasks used here targeted sentence comprehension, which encompasses a broad range of linguistic (e.g., lexical, combinatorial) functions. Both tasks also attempted to limit activation of unimodal sensory cortices (visual areas in READLOC, auditory areas in SPEECHLOC) as well as activity related to general task performance (e.g., task timing, attending to a screen, pressing buttons) through control conditions that employed similar stimuli and demands (Fedorenko et al. 2010; Scott et al. 2017). We were also able to assess the separability of transmodal from unimodal processing by focusing on the AUDLOC contrast. This map, which represented the contrast of listening to incomprehensible speech sounds against a passive fixation baseline, revealed robust bilateral activation within or near auditory cortex (Fig. 4, right; Rauschecker & Scott 2009; Humphries et al, 2010; Dick et al., 2017; Formisano et al., 2003), supporting that it was an effective activator of auditory areas. These activations also aligned with the iFC-defined AUD network (Fig. 4; left). Thus, despite being defined using independent task data (REST vs. SPEECHLOC) and different approaches (iFC vs. task activation), the boundaries of the LANG network nicely demarcated regions likely serving auditory functions (i.e., sounds that did not contain comprehensible words), and separated them from regions serving transmodal language functions (i.e., active for both SPEECHLOC and READLOC).

Further support for this demarcation was found by considering the overlap between the AUDLOC and READLOC task contrast maps (Fig. 5). We found that the boundaries of the LANG network encapsulated regions active during reading (Braga et al., 2020), and that generally these reading-related regions were not activated by the distorted speech clips. This was informative because the maps were defined in independent data: the distorted speech condition was not used as a control for the reading task, and therefore their lack of overlap was not dictated by the analysis strategy. Notably, the AUDLOC contrast map overlapped with regions that were activated by the SPEECHLOC contrast but which did not overlap with the READLOC contrast (and were outside of the LANG network). This separation suggests that the SPEECHLOC map included both auditory regions and transmodal regions (despite the subtraction of the control condition), which were well-separated by the iFC maps and the comparison of READLOC and AUDLOC maps (Fig. 5). These overarching findings bolster the claim that the LANG network is transmodal for language, despite being closely juxtaposed with regions supporting unimodal auditory functions.

Although the auditory localizer stimuli were successful in activating auditory cortex, the distorted speech stimuli still included recognizable aspects of speech. The filtering procedure muffled words to make comprehension difficult while preserving the intonation and prosody of each speaker (following Scott et al., 2017). Thus, the AUDLOC contrast likely still preserved activation of speech-specialized regions, including those supporting suprasegmental features of speech. Notably, in most individuals the AUDLOC maps included frontal lobe regions, close to the ventral motor strip, that likely do not serve auditory processing. Thus it is likely that the AUDLOC map at least partially included non-auditory regions. Prior work has shown that a similar distribution of regions, encircling auditory cortex in the temporal lobe and including regions near precentral cortex, may serve articulatory or phonetic processes (de Heer et al. 2017; see also the ‘INT’ network in Braga et al. 2020). Potentially, such a network may link precentral regions controlling the orofacial musculature with temporal regions that encode speech-like sound features including prosody and intonation. Such features may also explain why the auditory localizer activity often extended into the language network. Importantly, this interpretation remains consistent with the idea that the LANG regions encode a transmodal, not purely auditory, function. These results imply that there may be additional heterogeneity within the LANG network, as mapped with iFC, that might support intonation- or prosody-level speech processing. Interestingly, the overlap with activity during reading (READLOC; Fig. 5) also implies that that these processes are recruited by either visual or auditory language stimuli, suggesting they are also transmodal to some degree. To confirm this supposition, future studies might use fully non-linguistic acoustic stimuli (e.g., synthesized tones, music) and higher resolution data to yield further insights into the separability of these functions.

### 4.4 Limitations

While all participants completed the same number of runs for each task, we excluded some runs because of head motion or inconsistent behavioral responses. As a result, task activity maps were built from different quantities of data depending on the participant. For READLOC, four participants had all 8 runs included, two participants had 7 runs, one participant (S7) had 5 runs, and one participant (S4) had 4 runs (∼19 minutes). This last participant predictably showed less robust language activity for this task compared to the other participants. Nevertheless, the study that originated this visual language task reported that ∼15-20 minutes was sufficient to identify the language network (Fedorenko et al. 2010). Performance was more consistent for SPEECHLOC, with five participants including all 8 runs, and the remaining three participants including 7 runs. Participant S4 demonstrated a more robust activity map for this task. Thus, additional data may have improved consistency of results across participants.

A further limitation is that neither task required participants to demonstrate language comprehension. In theory, both tasks could be performed by ignoring the stimuli and simply waiting to respond to the button-press cue. If this were the case, however, we might expect activations to be comparable during both the comprehensible and incomprehensible conditions, rather than revealing classic language regions (and overlapping with the LANG network). Prior work has shown that the passive version of the language localizer task used here (Fedorenko et al. 2010; Scott et al. 2017) yields very similar activation to a version which requires a button press for recognized words (Fedorenko et al. 2010; Diachek et al., 2020). Further work could assess the extent to which these factors influence the degree of overlap between task activation and the iFC-defined network.

## 5 Conclusions

We show that the iFC-defined distributed language network is recruited during processing of language from multiple input modalities, and therefore supports a transmodal language function. In addition, we demonstrate that the boundaries of this network are a relatively good localizer for the separation between unimodal auditory and transmodal language functions. Further work is needed to determine the exact nature of this separation, including how factors such as intonation and prosody affect the underlying regions. Our results suggest that iFC performed within individuals is a useful tool for understanding the complex interactions of cognitive functions underlying language abilities.

## 6 Data and Code Availability

All data needed to evaluate the conclusions in the paper are present in the paper. Data files can be provided upon request.

## 7 Author Contributions

J.J.S. & R.M.B. designed the study; J.J.S. & R.M.B. collected the data; J.J.S. and N.L.A. analyzed the data with input from R.M.B; J.J.S., N.L.A. & R.M.B wrote and edited the manuscript.

## 10 Acknowledgements

We would like to thank Ania M. Holubecki and Maya Lakshman for their assistance with sound calibration for the SPEECHLOC clips.

This research was also supported in part through the computational resources and staff contributions provided for the Quest high performance computing facility at Northwestern University which is jointly supported by the Office of the Provost, the Office for Research, and Northwestern University Information Technology, and the Center for Translational Imaging at Northwestern University.

